# CHIC: a short read aligner for pan-genomic references

**DOI:** 10.1101/178129

**Authors:** Daniel Valenzuela, Veli Mäkinen

## Abstract

Recently the topic of computational pan-genomics has gained increasing attention, and particularly the problem of moving from a single-reference paradigm to a pan-genomic one. Perhaps the simplest way to represent a pan-genome is to represent it as a set of sequences. While indexing highly repetitive collections has been intensively studied in the computer science community, the research has focused on efficient indexing and exact pattern patching, making most solutions not yet suitable to be used in bioinformatic analysis pipelines.

**Results:** We present CHIC, a short-read aligner that indexes very large and repetitive references using a hybrid technique that combines Lempel-Ziv compression with Burrows-Wheeler read aligners.

**Availability:** Our tool is open source and available online at https://gitlab.com/dvalenzu/CHIC

## 1 Introduction

Research in computational pan-genomics has gained increasing attention, specially since recent initiatives are curating huge repositories of human genomes [4, 3, 5], posing the challenge of how to handle those large repositories efficiently.

While there is no consensus on how to represent a pan-genomic reference [2], many bioinformatics research projects have studied pan-genomic read alignment using a specific structure like a graph or a reference plus variations [18, 10, 19, 6, 7, 14]. On the other hand, the simpler model of the pan-genome as a set of sequences has been mostly studied from a computer science perspective [15, 16, 8], where the focus has been on efficient indexing and exact pattern matching, but to the best of our knowledge, no off-the-shelf solution for read alignment has been provided.

Here we introduce CHIC aligner, an off-the-shelf read aligner that is capable of indexing very large and repetitive references to align short reads to them very efficiently.

CHIC is a generalized version of the exact pattern matching tool CHICO [20] which uses an Hybrid Index [8] that combines Lempel-Ziv compression techniques with Burrows-Wheeler based indexing.

## 2 Methods

### Index construction

CHIC’s first step to index the reference is to compute a LZ77-compatible parsing [20]. This stands for greedy or non-greedy LZ77 parsing [21], Relative Lempel-Ziv (RLZ) [11] or variants of these. Those algorithms, originally designed to compress a sequence, tokenize the reference into phrases of two types. The first one, a ‘copying phrase’, corresponds to a substrings that have a previous occurrences in the input, and therefore can be represented as a copy using a pair of integers indicating the previous occurrence in the text and the length of the repetition. The second type, a ‘literal phrase’, corresponds to a string that does not have a known previous occurrence and therefore must be represented explicitly when these algorithms are used to compress the input. As the text is more repetitive, the copying phrases became longer, which accounts for the compression effectiveness of the LZ algorithms.

CHIC’s second step uses the LZ parsing and the input reference to build the so called kernel sequence [9]. The idea of the kernel sequence is to remove most of the redundancy from the input sequence, preserving only crucial substrings. The kernel sequence is initially empty, and the text is scanned once from left to right. Each time a literal phrase is found, its content is entirely appended to the kernel sequence. Each time a copying phrase is found only the first and the last *M* characters of the phrase are appended to the kernel sequence, where *M* is a user-defined parameter that sets an upper bound on the length of the queries. The remaining characters of the copying phrases can be safely discarded because there is a previous occurrence of them that have been already added to the kernel sequence.

CHIC’s third step is to index the kernel sequence using a standard read aligner. By default, it uses Bowtie2 [12], but the user can indicate through a parameter to use BWA instead.

The fourth step is to encode the LZ parsing so that at query time the alignments and its repetitions can be reported efficiently. This includes an encoding of the phrase boundaries in the input reference and the corresponding projections into the kernel sequence, and a succinct representation of the copying phrases. **Read alignment.** First the reads are aligned into the kernel sequence using the designated read aligner (by default Bowtie2). Then those alignments are mapped back to the original reference sequence, using rank and select [17] queries on the phrase boundaries. These are the sole output in the default reporting level (primary occurrences).

The second level of reporting (maximum) considers that for each primary occurrence there might be identical copies were discarded when content from the copying phrases was discarded. Using the encoding of the copying phrases and the phrase boundaries CHIC can report all the identical copies of the primary occurrences.

CHIC’s two reporting levels can be combined with the reporting levels accepted by Bowtie2 (or BWA) in the kernel sequence. That is, the user can specify that he wants to retrieve the *k* best alignments from the kernel sequence, and whether or not he wants to retrieve the identical copies of them.

### Resource usage

Computing the LZ77 parsing can require significant resources and it can be a bottleneck to build the index, specially when the input reference is too big to fin in main memory. In this case, CHIC aligner can resort to RLZ algorithms, avoiding the use of external memory algorithms and leveraging parallelism to achieve a fast index construction.

### Input and output

Similar to standard read aligners, CHIC uses a reference sequence in multi-fasta format, reads in FASTQ format, and the alignments are output in the BAM format [13]. This makes our tool ready to use in sequencing pipelines.

### Experimental results

To demonstrate the practical performance of CHIC we build references using different number of human genomes from the 1000 genomes project.

The input reference sequence was built based on GRCh37 plus variation observed in a subset of individuals from the 1000 genomes project [4]. A subset of either 10 or 100 individuals was taken, each individual was represented as a diploid genome yielding in total 20 or 200 reference sequences on top of GRCh37.

To carry out our experiment in a realistic set we aligned reads from an individual that is not part of the pan-genome reference. We took a sample of 10^6^ paired-end reads from a recent study on the genome of a Mongolian individual [1]. We measured the indexing time for each pan-genomic reference, the size of the resulting index and the total time to align the reads to the reference. Also we report the number of mapped reads. As a baseline, we compare against Bowtie2 using a single human genome as a reference. Table 1 summarize the results.

**Table 1.** Scalability of CHIC aligner. Time required to build the index, to align a set of 10^6^ paired-end reads, and the number of unaligned reads. BOWTIE2_1_ indexes the human reference, CHIC_*x*_ indexes *x* different versions of the human genome (only numbered chromosomes).

| Aligner | Input size | Indexing |  | Alignment |  |
| --- | --- | --- | --- | --- | --- |
|  |  | Time | Index Size | Time | Mapped reads |
| BOWTIE2<sub>1</sub> | 2.7GB | 1h | 3.708GB | 5.32s | 782883 |
| CHIC<sub>21</sub> | 57GB | 5h | 25GB | 2m | 785552 |
| CHIC<sub>201</sub> | 540GB | 35h | 180GB | 24m | 786026 |

## 3 Discussion

Our results show that CHIC can effectively handle very large reference sequences and align reads to them. As expected, the use of a pan-genomic reference instead of a single reference results in an increased number of mapped reads.

An important open problem is how to define a meaningful scoring functions in the pan-genomic context. While in the traditional (single-reference) setup, an alignment that is not unique is not desirable, for pan-genomic references unique maps are much rare and non-unique alignments should not be discarded a priori, hence the need of a scoring function that is pan-genome- aware.

## Acknowledges

Many thanks to Esa Pitköanen for pointing out some recent references on se-quencing projects.

